# Megafaunal extinctions—not climate change—seem to explain Holocene genetic diversity declines in *Numenius* shorebirds

**DOI:** 10.1101/2021.07.02.450884

**Authors:** Hui Zhen Tan, Justin J.F.J. Jansen, Gary A. Allport, Kritika M. Garg, Balaji Chattopadhyay, Martin Irestedt, Sean E.H. Pang, Glen Chilton, Chyi Yin Gwee, Frank E. Rheindt

## Abstract

Understanding the relative contributions of historical and anthropogenic factors to declines in genetic diversity is important for informing conservation action. Using genome-wide DNA of fresh and historic specimens, including that of two species widely thought to be extinct, we investigated fluctuations in genetic diversity and present the first complete phylogenomic tree for all nine species of the threatened shorebird genus *Numenius*, known as whimbrels and curlews. Most species faced sharp declines in effective population size, a proxy for genetic diversity, soon after the Last Glacial Maximum (around 20,000 years ago). These declines occurred prior to the Anthropocene and in spite of an increase in breeding area predicted by environmental niche modelling, suggesting that they were not caused by climatic or recent anthropogenic factors. Crucially, these genetic diversity declines coincide with mass extinctions of mammalian megafauna in the Northern Hemisphere. Demise of ecosystem-engineering megafauna which maintained open habitats would have been detrimental for grassland and tundra-breeding *Numenius* shorebirds. Our work suggests that the impact of historical factors such as megafaunal extinction may have had wider repercussions on present-day population dynamics of open habitat biota than previously appreciated.

## Introduction

Rates of population decline and extinction have risen sharply during the ongoing sixth mass extinction crisis (Ceballos, Ehrlich, & Raven, 2020; Dirzo & Raven, 2003; Sánchez-Bayo & Wyckhuys, 2019; S. N. Stuart et al., 2004). Species distribution models based on future climate scenarios forecast that rates of endangerment will further accelerate, underscoring the need for conservation action (Thomas et al., 2004). In this era of increasing biodiversity loss, the maintenance of genetic diversity within species has become a focus of conservation as it is thought to predict evolutionary adaptability and extinction risk (Frankham, 2005; Hoban et al., 2020; Jetz et al., 2014). Modern declines in genetic diversity have been documented for a handful of species (Allentoft & O’Brien, 2010; Chattopadhyay, Garg, Mendenhall, & Rheindt, 2019; S. R. Evans & Sheldon, 2008; Garner, Rachlow, & Hicks, 2005), but we continue to know little about the global mechanisms of genetic diversity loss.

Anthropogenic climate change is widely recognised for its pervasive impact on biodiversity and genetic diversity (C. N. Johnson et al., 2017; Miraldo et al., 2016; Turvey & Crees, 2019). However, historical events have equally left their signature in the genetic profiles of present-day species (Hewitt, 2000). Comparative genomics of extinct versus extant species could add an important perspective to elucidating such trends in faunal endangerment (Frankham, 2005).

We used a museomic approach to investigate fluctuations in effective population size in all nine species of the migratory shorebird genus *Numenius*, known as whimbrels and curlews, including two species, the slender-billed curlew (*N. tenuirostris*) and Eskimo curlew (*N. borealis*), that are presumed to be extinct (Buchanan et al., 2018; Butchart et al., 2018; Kirwan, Porter, & Scott, 2015; Pearce-Higgins et al., 2017; Roberts, Elphick, & Reed, 2010; Roberts & Jarić, 2016). Members of the genus *Numenius* breed across the Northern Hemisphere’s tundras and temperate grasslands, and are particularly vulnerable to endangerment due to comparatively long generation times (Pearce-Higgins et al., 2017).

Our objective was to characterise genetic diversity fluctuations in *Numenius* shorebirds, assess the relative impact of historical and anthropogenic factors on these fluctuations, and determine the mechanisms that may have had the biggest impact on their populations. Because of their dependence on open habitats, we expected the genetic diversity trends of whimbrels and curlews to track the availability of such habitats across the late Quaternary. We also expected significant declines in genetic diversity during the late Holocene when global human activity intensified, not least because the demise of the two extinct species has been attributed to habitat loss and hunting (Committee on the Status of Endangered Wildlife in Canada, 2009; Gallo-Orsi & Boere, 2001). By testing the timing of genetic diversity fluctuations against that of important ecological events, we could home in on the factors that influenced the evolutionary trajectory of this threatened shorebird lineage over the last ~20,000 years.

## Results and discussion

We sequenced 67 ancient and fresh samples across all nine *Numenius* species for target enrichment (Figure 1A; Table S1). After filtering for quality, a final alignment of 514,771bp across 524 sequence loci was retained for each of 62 remaining samples at a mean coverage of 118X. Phylogenomic analyses using MP-EST (L. Liu, Yu, & Edwards, 2010) revealed two separate groups, here called the “whimbrel clade” and the “curlew clade”, that diverged approximately 5 million years ago (Figure 1B; Figure S1A). This is the first phylogenomic tree to include all members of the genus *Numenius*. The use of degraded DNA from toepads of museum specimens allowed us to include the two presumably extinct taxa. Of these, the slender-billed curlew emerged as sister to the Eurasian curlew (*N. arquata*), a phenotypically similar species that occurs in sympatry in Central Asia (Sharko et al., 2019). On the other hand, the Eskimo curlew emerged as a distinct member of the curlew clade with no close relatives (Figure 1B). Our phylogenomic dating analyses demonstrated that 40.6% of the evolutionary distinctness (Jetz et al., 2014) of the curlew clade has been lost with the presumable extinction of the two species, and that another 15% is endangered (Figure 1B; Table S2).

**Figure 1.**
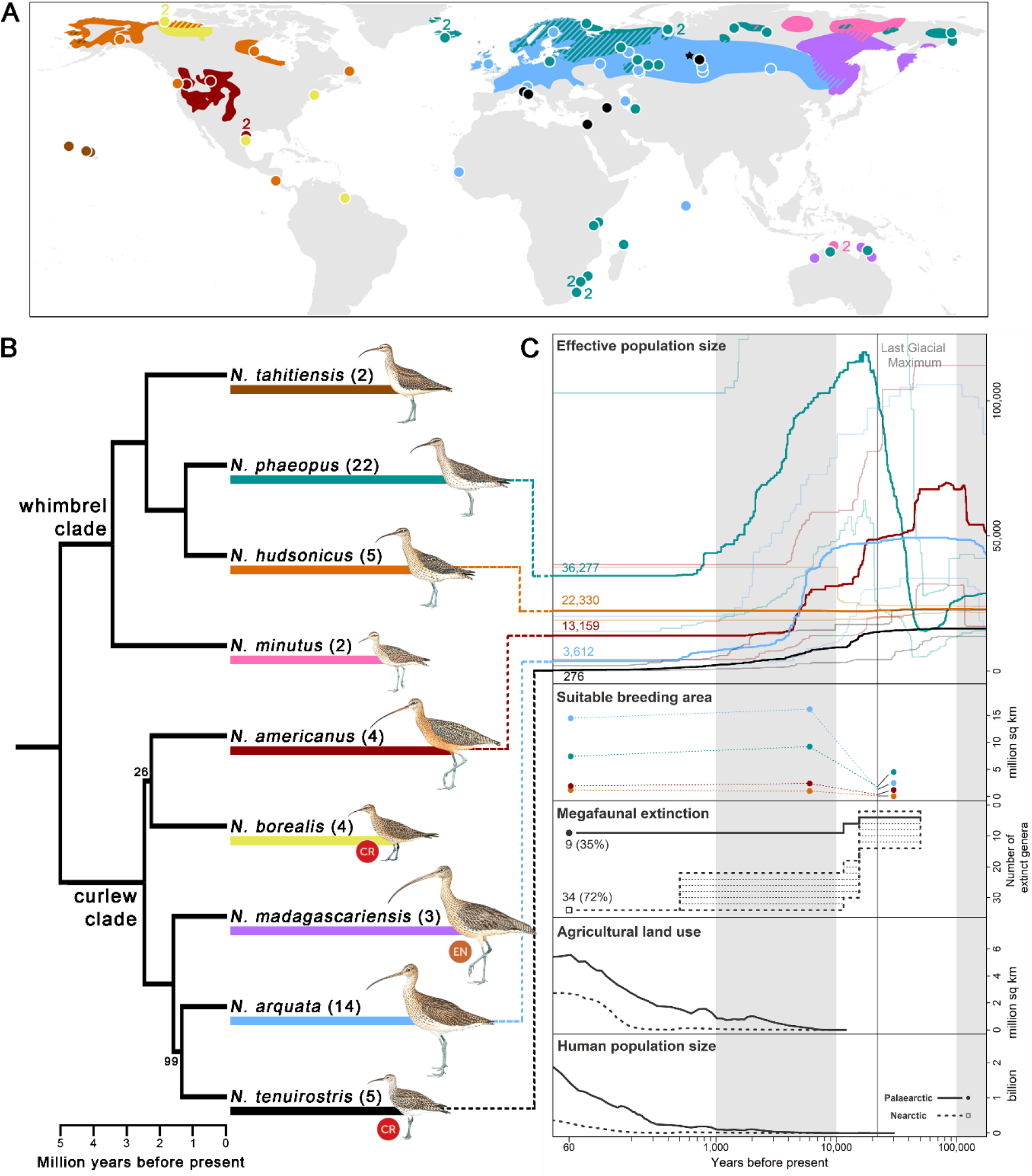
(A) Breeding distribution map and sampling localities of each *Numenius* species (BirdLife International and Handbook of the Birds of the World, 2017; Lappo, Tomkovich, & Syroeckovskiy, 2012); wintering and migratory ranges are not shown. Colours correspond to species identities in (B). Diagonal lines denote regions with co-distributed species. Each circle represents one sample unless otherwise specified by an adjacent number. The only known breeding records of *N. tenuirostris* were from near the village of Krasnoperova c.10 km south of Tara, Omsk (Russia), which is denoted by a black star (★), although this might not have been the core breeding area. (B) Phylogenomic tree constructed from an alignment of 514,771bp across 524 sequence loci. Tree topology (including bootstrap support values) and divergence times were estimated with MP-EST and MCMCTree respectively. Only bootstrap <100 is displayed. Sample sizes for each species are given in brackets. IUCN Red List status of critically endangered (CR) and endangered (EN) species are indicated. Illustrations of *Numenius* birds were reproduced with permission from Lynx Edicions. (C) Results of demographic history reconstruction using stairway plot for selected species displayed with key climatic, biotic and anthropogenic events. Effective population size: Line colours correspond to species identities in the tree in (B) and numbers at present time represent present-day effective population sizes. Thick lines represent the median effective population size while thin lines represent the 2.5 and 97.5 percentile estimation. The vertical grey line denotes the Last Glacial Maximum (LGM) and panels are shaded to aid reference to the time axis. Suitable breeding area: predicted suitable breeding area at LGM (22,000 years ago), mid-Holocene (6,000 years ago) and present day (1960–1990) using Maxent. Dot colours correspond to species identities in the tree in (B). Dotted lines connecting the dots are for visualisation purposes and do not represent fluctuations in breeding area. The following panels display the timings of key climatic, biotic and anthropogenic events including megafaunal extinction (in terms of number of extinct genera with dotted shading denoting uncertainty in estimates; Koch & Barnosky, 2006),_agricultural land use and human population size (HYDE 3.2; Goldewijk, Beusen, Doelman, & Stehfest, 2017; Goldewijk, Beusen, & Janssen, 2010). Line type corresponds to geographical area (Nearctic versus Palaearctic) as denoted in the ‘Human population size’ panel.

To characterise the differential impacts of extinction pressures, we reconstructed the demographic history of *Numenius* shorebirds. For five species with a sufficiently high sample size, we employed stairway plots (X. Liu & Fu, 2020) to infer fluctuations in effective population size (*N_e_*), a proxy for genetic diversity, given that this method works well for reduced representation genomic datasets such as ours, and has a relatively high accuracy for reconstructions of diversity changes in the Late Quaternary (X. Liu & Fu, 2020). Fluctuations in *N_e_* were compared against key biotic and anthropogenic events of the Late Quaternary. We also accounted for climatic changes by modelling the extent of suitable breeding areas of each species under present day (1960–1990), mid-Holocene (6,000 years ago), and LGM (22,000 years ago) climate conditions using the MaxEnt algorithm (Phillips, Anderson, & Schapire, 2006).

The Last Glacial Period preceding the LGM saw ice sheets at their maximum extent (Hughes, Gibbard, & Ehlers, 2013). During this time, tundra habitats dominated the northern latitudes and an increase in *N_e_* in the tundra-inhabiting Eurasian whimbrel (*N. phaeopus*) was observed (Binney et al., 2017; Y. Wang et al., 2021; Zimov et al., 1995; Figure 1C). Soon after, during the Pleistocene-Holocene transition, our stairway plots revealed generally sharp declines of *N_e_* in most species despite an increase in area of suitable breeding habitat predicted (Figure 1C). The extent of breeding habitat predicted by our ecological niche models relied on bioclimatic variables, suggesting that favourable conditions for *Numenius* shorebirds in the lead-up to the Holocene did not trigger an increase in genetic diversity, but instead coincided with precipitous declines in *N_e_*. To understand the drivers of *N_e_* declines in *Numenius* shorebirds, factors other than climate change would need to be considered.

During the Pleistocene-Holocene transition (starting at roughly 20,000 years ago), a mass extinction of megafaunal mammals (≥ 44 kg) was underway, known as the Late Quaternary Extinctions (Hedberg, Lyons, & Smith, 2022; C. N. Johnson, 2009), with most becoming extinct by 10kya (Figure 1C) (Koch & Barnosky, 2006; A. J. Stuart, 2015). Megafaunal mammals are ecosystem-engineers that maintain open landscapes such as temperate grasslands and steppes through grazing, browsing and physical impacts (Bakker et al., 2016; Goheen et al., 2018). During the intervening period between their extinction and the spread of ungulate domestication, there would have been no functional replacements for these ecosystem services (Hedberg et al., 2022; Lundgren et al., 2020). Open grasslands would have been subject to increasing forest succession (C. N. Johnson, 2009) and the amount of suitable habitat for *Numenius* shorebirds might have been less than predicted by forecasts relying only on bioclimatic variables. Therefore, genetic diversity fluctuations in *Numenius* shorebirds run counter to expectations based on natural climate change and seem to be best explained by the demise of the ecosystem-engineers that would have helped maintain shorebird breeding habitats.

By the late Holocene, the genetic diversity of most *Numenius* shorebirds stabilised at a time when anthropogenic impact was only starting to expand across the Northern Hemisphere with a steep rise in human population and land conversion for agriculture (Figure 1C). The timing of these events is inconsistent with the hypothesis that direct anthropogenic activity has been the main cause of genetic diversity declines in *Numenius* (Crisp, Trewick, & Cook, 2011). Events unrelated to modern anthropogenic pressure seem to have played a bigger role in the declines observed in *Numenius* shorebirds (Lucena-Perez et al., 2020; Nadachowska-Brzyska, Li, Smeds, Zhang, & Ellegren, 2015). It is possible that additional adverse effects caused by more recent anthropogenic impacts are not yet reflected in the genomes investigated, perhaps exacerbated by shorebirds’ long generation times.

At present, members of the curlew clade, which predominantly breed in temperate grasslands at lower latitudes, generally exhibit levels of *N_e_* that are lower than those of the higher latitude-breeding whimbrels (Figure 1C). Temperate grasslands face far greater anthropogenic pressures from land use than the northerly tundra (Pimm et al., 2014), contributing to further declines in curlews more so than in whimbrels. Genetic diversity estimates were lowest in the presumably extinct slender-billed curlew *N. tenuirostris* (Figure 1C). Low genetic diversity may contribute to a species’ extinction risk (Frankham, 2005; Spielman, Brook, & Frankham, 2004), although such links must be examined for each species independently and could possibly be conflated with other factors such as total population size (S. R. Evans & Sheldon, 2008; Teixeira & Huber, 2021).

Our study uncovers substantial declines in genetic diversity in curlews and whimbrels across the Late Quaternary. Analysing *N_e_* fluctuations over time allowed us to test which factors may have coincided with genetic diversity declines. Of the factors investigated, megafaunal extinctions—not natural climate change in the post-glacial period—best explain these declines and may have had cascading effects on species’ evolutionary trajectories that continue to impact them to the present day. Our results underscore that grassland biomes and their biota face unique challenges that warrant more conservation attention (Ceballos et al., 2010; Chan, Lacey, Pearson, & Hadly, 2005; Helm et al., 2009; Nakahama, Uchida, Ushimaru, & Isagi, 2018; Török, Ambarlı, Kamp, Wesche, & Dengler, 2016; Wesche et al., 2016). Our work demonstrates that relatively brief evolutionary events, such as the Late Quaternary Extinctions of megafauna, can have long-lasting evolutionary effects on populations, in our case for roughly ~10,000 years. The plight of *Numenius* shorebirds is a sobering reminder of the importance of conserving remaining genetic diversity to ensure the resilience of our planet’s biota.

## Materials and methods

### Taxon sampling

We acquired samples for all nine species in the genus *Numenius*, encompassing most of the known subspecies. Species and subspecies identities are as provided by the source museum or institution (Table S1) or assigned in reference to known breeding and wintering locations (Birds of the World, 2022). We also included one common redshank *Tringa totanus* as an outgroup for phylogenetic rooting. All samples were acquired through museum loans except for an individual of the endangered subspecies *N. phaeopus alboaxillaris* that was sampled during fieldwork by GAA (Table S1). Where possible, we acquired fresh samples (tissue or blood) because of their higher DNA quality. To represent rarely-sampled or presently-rare taxa for which no fresh samples were available, we acquired toepad material from historic museum specimens and applied ancient DNA methods.

### Baits design for target capture

We used the *Calidris pugnax* genome (accession no. GCA_001458055.1) (Küpper et al., 2015) to design baits to capture selected exons. We used EvolMarkers (C. Li, Riethoven, & Naylor, 2012) to identify single-copy exons conserved between *C. pugnax, Taeniopygia guttata* (accession no. GCF_003957565.1; released by the Vertebrate Genomes Project) and *Ficedula albicollis* (accession no. GCA_000247815.1). Exons longer than 500 bp with a minimum identity of 55% and an e-value < 10*e*^-15^ were isolated with bedtools 2.28.0 (Quinlan & Hall, 2010), forming our target loci. Only target loci with 40–60% GC content were retained and any overlapping loci were merged (Quinlan & Hall, 2010). Target loci with repeat elements were then filtered out in RepeatMasker 4.0.6 (Smit, Hubley, & Green, 2015). We arrived at a final set of 565 unique target loci with a mean length of 970bp. These target loci were used to design 19,003 100 bp-long biotinylated RNA baits at 4X tiling density (MYcoarray/Arbor Biosciences, USA).

### Laboratory methods

Both fresh and historic samples were subjected to DNA extraction, followed by library preparation and target enrichment, with slight modifications for various sample types to optimise yield. DNA extractions of fresh samples were performed using the DNEasy Blood & Tissue Kit (Qiagen, Germany) with an additional incubation step with heat-treated RNase. Extractions for historic samples were performed using the same kit but with modifications (Chattopadhyay et al., 2019). Historic samples were washed with nuclease-free molecular grade water before extraction and dithiothreitol was added to the digestion mix. DNA precipitation was performed for at least 12 hours and MinElute Spin Columns were used for elution (Qiagen, Germany). Historic samples were processed in a dedicated facility for highly degraded specimens.

DNA extracted from fresh samples was sheared via sonification using Bioruptor Pico (Diagenode, Belgium) to a target size of 250 bp. DNA extracts from historic samples were generally smaller than the target size; hence no further shearing was performed. Whole-genome libraries were prepared using the NEBNext Ultra™ II DNA Library Prep Kit for Illumina (New England Biolabs, Ipswich, USA) with modifications for subsequent target enrichment. For fresh samples, adaptor concentrations were kept constant regardless of DNA input amount. Size selection with AMPure XP beads (Beckman Coulter, USA) was performed for 250 bp insert sizes. The reaction was split into two equal parts before polymerase chain reaction (PCR) amplification and combined afterwards for subsequent steps. For historic samples, a formalin-fixed, paraffin-embedded (FFPE) DNA repair step was first performed using NEBNext FFPE DNA Repair Mix (New England BioLabs). A 10-fold dilution of adaptors was used, and no size selection was performed. For both types of samples, twelve cycles of PCR amplification were performed.

Target enrichment was carried out following the MYbaits manual (Arbor Biosciences, USA) with modifications (Chattopadhyay et al., 2019). We used 1.1 uL of baits per fresh sample (~5X dilution) and 2.46 uL of baits per ancient sample (~2X dilution). For fresh samples, hybridization of baits and target loci was performed at 65°C for 20 hours and 15 cycles of amplification were performed. For historic samples, hybridization was performed at 60°C for 40 hours, and 20 cycles of amplification were performed. For both fresh and historic samples, one negative control sample was added for each batch of extraction, library preparation and target enrichment. Extracts, whole-genome libraries, final enriched libraries, and all negatives were checked for DNA concentration on a Qubit 2.0 Fluorometer using the Qubit dsDNA HS assay kit (ThermoFisher Scientific, USA), and for fragment size on a Fragment Analyzer using the HS NGS Fragment kit (1–6000 bp) (Agilent Technologies Inc., USA). Final enriched libraries were pooled at equimolar quantities. A total of 67 enriched libraries were sequenced, with fresh and historic samples sequenced separately on two Illumina HiSeq 150 bp paired-end lanes (NovogeneAIT, Singapore).

### Reference genome assembly

We obtained a sample of *N. phaeopus* (ZMUC 112728) from the Natural History Museum of Denmark, Copenhagen, for reference genome assembly. Its genomic DNA was extracted using the KingFisher™ Duo Prime Magnetic Particle Processor (ThermoFisher Scientific, USA) and the KingFisher Cell and Tissue DNA Kit (Thermo Fisher Scientific). A linked-read sequencing library was prepared using the Chromium Genome library kits (10X Genomics) and sequenced on one Illumina Hiseq X lane at SciLifeLab Stockholm (Sweden). The *de novo* assembly analysis was performed using 10X Chromium Supernova (v. 2.1.1). Reads were filtered for low quality and duplication, while assemblies were checked for accuracy and coverage and the best assembly was selected based on the highest genome coverage with the fewest errors. The final genome had a size of 1.12 Gb at a coverage of 50X with N50 = 3504.2kbp.

### Raw reads processing

Raw reads were checked for sequence quality in FastQC 0.11.8 (Babraham Bioinformatics) and trimmed to remove low-quality termini and adaptors in fastp 0.20.0 (Chen, Zhou, Chen, & Gu, 2018). We retained reads with a minimum length of 36bp and set a phred quality threshold of 20. Retained reads started at the first base satisfying minimum quality criteria at the 5’-end and were truncated wherever the average quality fell below the threshold in a sliding window of 5bp. Duplicates were removed using FastUniq 1.1 (Xu et al., 2012) before sequence quality, duplication rate and adaptor content were checked again in FastQC. We employed FastQ Screen 0.14.0 (Wingett & Andrews, 2018) to assign the source of DNA against a list of potential contaminants. We aligned reads to our assembled *Numenius phaeopus* genome, *Homo sapiens* (accession no. GCF_000001405.39), and a concatenated database of all bacterial genomes available on GenBank (National Center for Biotechnology Information (NCBI), 1988). Only reads that mapped uniquely to the *N. phaeopus* genome were retained. Reads were sorted and re-paired using BBtools 37.96 (Bushnell, 2014). Downstream bioinformatic procedures were split into single nucleotide polymorphism (SNP)-based and sequence-based analyses.

### SNP calling

For SNP-based analyses, reads were aligned to the target sequences used for bait design with bwa-mem 0.7.17 (Li, 2013). The output alignment files were converted to bam files (view) and sorted by coordinates (sort) using SAMtools 1.9 (H. Li et al., 2009). Alignments were processed in Picard 2.20.0 (Picard tools, Broad Institute, Massachusetts, USA) to add read group information (AddOrReplaceReadGroups), and another round of duplicate identification was performed (MarkDuplicates) before alignment files were indexed (BuildBamIndex). The reference file of target sequences was indexed in SAMtools (faidx) and a sequence dictionary was created in Picard (CreateSequenceDictionary). To improve SNP calling accuracy, indel realignment was performed in GATK 3.8 (McKenna et al., 2010) (RealignerTargetCreator, IndelRealigner). We inspected historic DNA alignments in mapDamage 2.0.9 (Jónsson, Ginolhac, Schubert, Johnson, & Orlando, 2013) and trimmed up to 5bp from the 3’ ends of both reads to minimise frequencies of G to A misincorporation (<0.1) and soft clipping (<0.2). Finally, alignments were checked for quality and coverage in QualiMap 2.2.1 (Okonechnikov, Conesa, & García-Alcalde, 2016).

We first generated likelihoods for alignment files in BCFtools 1.9 (Li, 2011) (mpileup), skipping indels. Using the same program, we then called SNPs (call) for all *Numenius* samples using the multiallelic and rare-variant calling model. Called SNPs were filtered in VCFtools 0.1.16 (Danecek et al., 2011) to retain sites with quality values >30, mean depth 30–150, minor allele frequency ≥0.02 and missing data <5%, in this order. Missingness and depth of sites and individuals, respectively, were quantified for SNPs called. We removed eight individuals from downstream analyses due to a combination of high missing data (>0.4%) and low coverage (<36X), yielding a SNP set representing 58 samples. A Perl script (rand_var_per_chr.pl) was used to call one SNP per locus to avoid calling linked SNPs (Caballero, 2018). SNPs were further screened for linkage disequilibrium in PLINK 1.9 (Purcell et al., 2007) using a sliding window of 50 SNPs with a step size of 10 and an r^2^ correlation threshold of 0.9. We also screened for neutrality of SNPs in BayeScan 2.1 (Foll & Gaggiotti, 2008) using default settings. We additionally created a dedicated SNP set per species for input into demographic history reconstruction using the method described above, but without minor allele frequency cut-offs and with all SNPs at each locus retained.

### Population genomic analyses

We conducted principal component analysis (PCA) for all *Numenius* samples using the R package SNPRelate 1.16.0 (R Core Team, 2022; Zheng et al., 2012) (Figure S1A). We did not detect any considerable genomic differentiation along subspecific delimitations within *N. phaeopus* and *N. arquata*, whose population-genetic structure had been resolved with thousands of genome-wide markers in a previous study (Tan et al., 2019) (Figure S1B, C). Samples of *N. p. alboaxillaris* and *N. a. suschkini*, two Central Asian taxa that are described in the literature as phenotypically differentiated (Allport, 2017; Engelmoer & Roselaar, 1998b, 1998a; Morozov, 2000), did not emerge as genomically distinct from other conspecific populations and are likely to represent ecomorphological adaptations controlled by few genes. Sample NBME 1039630, which had been labeled as *N. tenuirostris*, and sample MCZR 15733, which was initially identified as an *N. arquata* that shares many morphological features with *N. tenuirostris*, clustered with *N. arquata* samples (Table S1, Figure S1D). Both samples were assigned to *N. arquata* in subsequent phylogenetic analyses.

### Sequence assembly

For sequence-based analyses, reads were assembled using HybPiper 1.3.1 (M. G. Johnson et al., 2016) (reads_first) to yield sequence loci. Firstly, reads were mapped to the target sequences using BWA 0.7.17 (H. Li & Durbin, 2009) and sorted by gene. Contigs were then assembled from the reads mapped to respective loci using SPAdes 3.13 (Bankevich et al., 2012) with a coverage cutoff value of 20. Using Exonerate 2.4.0 (Slater & Birney, 2005), these contigs were then aligned to the target sequences and sorted before one contig per locus was chosen to yield the final sequences. We inspected locus lengths (get_seq_lengths) and recovery efficiency (hybpiper_stats) across all loci. We then investigated potentially paralogous loci (paralog_investigator) by building gene trees using FastTree 2.1.11 (Price, Dehal, & Arkin, 2010) (paralog_retriever), leading to the removal of 10 loci. All loci retained were present in at least 80% of individuals and constituted at least 60% of the length of total target loci. In summary, a total of 525 loci with a mean length of 969 bp (492–6,054 bp) were recovered from 62 samples.

### Phylogenomic analyses using sequence data

Multisequence alignment was performed for each locus using MAFFT 7.470 (Katoh & Standley, 2013), allowing for reverse complement sequences as necessary. Alignments were checked for gaps using a custom script, and loci with >35% gaps were removed from downstream analyses. A total alignment length of 514,771 bp was obtained.

Phylogenomic analyses were performed on a concatenated dataset as well as on individual gene trees. Concatenation was performed with abioscript 0.9.4 (Larsson, 2010) (seqConCat). For the concatenated dataset, we constructed maximum-likelihood (ML) trees using RAxML 8.2.12 (Stamatakis, 2014) with 100 alternative runs on distinct starting trees. We applied the general time reversible substitution model with gamma distributed rate variation among sites and with estimation of proportion of invariable sites (GTR+I+G) (Abadi, Azouri, Pupko, & Mayrose, 2019; Arenas, 2015).

For individual gene trees, the best substitution model for each locus was determined using jModelTest 2.1.10 (Darriba, Taboada, Doallo, & Posada, 2012) by virtue of the corrected Akaike information criterion value. We then constructed ML trees in PhyML 3.1 with the subtree pruning and regrafting algorithm, using 20 initial random trees. We performed 100 bootstrap replicates with ML estimates for both proportion of invariable sites and value of the gamma shape parameter. Individual gene trees were then rooted with Newick Utilities 1.3.0 (Junier & Zdobnov, 2010). We removed one locus from downstream analyses due to the absence of an outgroup sequence such that 524 loci were retained across 62 samples.

Species tree analyses were performed using the rooted gene trees in MP-EST 1.6 (L. Liu et al., 2010), without calculation of triple distance among trees. We grouped samples by species and performed three runs of 10 independent tree searches per dataset (Cloutier et al., 2019). To calculate bootstrap values of the species tree, we performed multi-locus, site-only resampling (Mirarab, 2014) from the bootstrap trees’ (100 per gene) output from PhyML. The resulting 100 files, each with 100 bootstrap trees, were rooted and species tree analyses were performed in the same manner for each file in MP-EST. The best tree from each run was identified by the best ML score and compiled. Finally, we used the majority rule in PHYLIP 3.695 (Felsenstein, 2009) to count the number of times a group descending from each node occurred so as to derive the bootstrap value (consense).

For estimation of divergence times, we applied MCMCtree and BASEML (dos Reis & Yang, 2011), a package in PAML 4.9e (Yang, 2007). To prepare the molecular data from 62 samples and 524 loci, we compiled the DNA sequence of each sample and combined all samples onto separate rows of the same file. We then obtained consensus sequences for each species using Geneious Prime 2020.2 (Kearse et al., 2012), with a majority support threshold of 50% and ignoring gaps. We visually checked the resulting consensus sequences to ensure that ambiguous bases remained infrequent. Consensus sequences were organised by loci as per the input format for MCMCtree. We then prepared the input phylogenetic tree using the topology estimated in MP-EST with calibrations of the two most basal nodes, namely between our outgroup (*Tringa totanus*) and all *Numenius* species, as well as that between the whimbrel and curlew clades within *Numenius*. Due to a lack of known fossils within the genus *Numenius*, we were unable to perform fossil node calibrations. Instead, we utilised p-distance values calculated from the COI sequences of *Numenius* species. Specifically, we applied the bird COI mutation rate of 1.8% per million years (Lavinia, Kerr, Tubaro, Hebert, & Lijtmaer, 2016) and converted mean, maximum and minimum p-distance values of both nodes to time (100 million years ago (MYA)). We maintained a conservative position and scaled the COI-based timings by a factor of two to obtain the final lower and upper bounds of node timings. We used the default probability of 0.025 that the true node age is outside the calibration provided.

To run MCMCtree, we first calculated the gradient and Hessian matrix of the branch lengths with the GTR substitution model applied, using default values of gamma rates and numbers of categories (mcmctree-outBV.ctl). We then performed two independent Markov chain Monte Carlo (MCMC) samplings of the posterior distribution of divergence times and rates (mcmctree.ctl). All default values were used except that a constraint on the root age was set to <0.3 (100 MYA). We also varied the prior for the birth-death process with species sampling and ensured that time estimates are not affected by the priors applied (dos Reis & Yang, 2019). We then performed convergence diagnostics for both runs in R to ensure that posterior means are similar among multiple runs, while checking that the parameter space has been explored thoroughly by the MCMC chain. Finally, we conducted MCMC sampling from the prior with no data to check the validity of priors used by comparing them with posterior times estimated. Again, two independent MCMC samplings were performed with convergence diagnostics.

Phylogenetic trees were visualised in FigTree 1.4.4 (Rambaut, 2018) with bootstrap values and node ages (MYA) including the 95% credibility intervals. Evolutionary distinctness and phylogenetic diversity was calculated for each branch (Jetz et al., 2014) using the divergence times estimated in MCMCTree.

### Demographic history reconstruction

We derived trends in effective population size using stairway plot 2.1.1, which uses the SNP frequency spectrum and is suitable for reduced representation datasets (X. Liu & Fu, 2020; Patton et al., 2019). From the dedicated SNP sets that were created without minor allele frequency cut-off, we calculated a folded site frequency spectrum using vcf2sfs.py 1.1 (Marques, Lucek, Sousa, Excoffier, & Seehausen, 2019). We assumed a mutation rate per site per generation of 8.11*e*^-8^, as estimated for shorebirds in the same order as *Numenius* (Charadriiformes) (X. Wang et al., 2019), and applied the following generation times respectively: *N. americanus* 7 years, *N. arquata* 10 years, *N. hudsonicus* 6 years, *N. phaeopus* 6 years, *N. tenuirostris* 5 years (Bird et al., 2020; IUCN, 2020). We ran stairway plot on all species, applying the recommended parameters.

Stairway plot is expected to perform at its highest accuracy in the reconstruction of demographic history in the recent rather than distant past. However, the definition of recent past varies from anywhere between 30 generations to ~40,000 generations before present (X. Liu & Fu, 2015, 2020; Patton et al., 2019). We did not set a cutoff for the time period investigated but let it be determined by the program itself. Additionally, we omitted reconstructions of the last 10 steps to avoid overinterpretation of the distant past (X. Liu & Fu, 2015). We only displayed the results from the time period for which there was data across all species, and only for four species represented by five or more samples (stairway_plot_es Stairbuilder), as recommended for accurate inference (X. Liu, personal communication, October 14, 2020). We later also included *N. americanus*, for which we had four samples, as its sample size did not appear to affect the reliability of results (Figure 1). We were unable to include the remaining species (*N. borealis, N. tahitiensis*, and *N. minutus*) as their demographic history reconstructions were clearly affected by a lack of sufficient sample size. For *N. borealis*, two out of the five samples showed high missingness, with adverse effects on stairway plot analyses, both in runs including all five samples and those that excluded the two samples of high missingness (Figure S2). Our ability to trial a large number of samples for laboratory work was also limited by the availability of target enrichment baits.

We attempted to infer demographic history using sequentially Markovian coalescent-based methods, which are more reliable for older timescales, to corroborate our stairway plot results (Patton et al., 2019). In particular, we used the Pairwise Sequentially Markovian Coalescent (PSMC) model (H. Li & Durbin, 2011) as it has been successfully applied to reduced-representation datasets (S. Liu & Hansen, 2017). This method allows for analyses of all species as only one sample per species is required as input. However, given the constraints created by the sampling density of our target enrichment dataset, we were unable to run PSMC successfully.

### Ecological Niche Modelling

We performed ecological niche modelling (Peterson et al., 2011) to predict the extent of suitable breeding area for species across the duration of our demographic history reconstruction. We were able to do so for each species in the stairway plot except *Numenius tenuirostris* due to the paucity of confirmed breeding records. We obtained species occurrence data from eBird (2021) and the Global Biodiversity Information Facility (GBIF; using only records with coordinate uncertainty < 1,000 m) (GBIF.org, 2022a, 2022b, 2022c, 2022d, 2022e, 2022f). For *N. phaeopus*, we also included confirmed breeding localities from Lappo et al. (2012) to improve sample size. Species occurrence data from various sources were combined and further filtered (Table S3). Occurrence points were filtered by month to retain only records in peak breeding months of respective species (Birds of the World, 2022). For species with sufficient occurrence points, occurrence points were also filtered by year to match the time range of the climatic variables, i.e., 1960–1990. Otherwise, occurrence records from all years were used to maximise sample size. For species that span the entire Palaearctic (*N. phaeopus* and *N. arquata*), sampling density was much higher in Europe. To account for the extreme sampling bias, in addition to generating a kernel density estimate (see next paragraph), occurrence records within Europe for these two species were randomly down-sampled to match sampling density across the rest of the Palearctic. Occurrence records outside of the known breeding area of each species were removed (BirdLife International and Handbook of the Birds of the World, 2017; Lappo et al., 2012). Finally, to reduce spatial autocorrelation, occurrence records were thinned using a 50 km buffer (Aiello-Lammens, Boria, Radosavljevic, Vilela, & Anderson, 2015).

To account for sampling bias specific to shorebirds, such as those of this study, we generated a kernel density estimate using the R package spatialEco 1.3-7 (J. Evans, 2021) based on the occurrences of species within Scolopacidae. The kernel density estimates was then used to inform background point selection (i.e., matching sampling bias) (Kramer-Schadt et al., 2013). For each species, we further limited the sampling of background points to areas outside a 10 km buffer around occurrence points and within a 500 km buffer around the known breeding area using the R packages terra 1.5-21 and raster 3.5-15 (Hijmans, 2022b, 2022a). A total of 10,000 background points were then sampled without replacement for each species.

All 19 bioclimatic variables (raster; 2.5 arcmin resolution of ~4.5 km) from WorldClim 1.4 (Hijmans, Cameron, Parra, Jones, & Jarvis, 2005) were obtained for the present day (1960–1990), mid-Holocene (6,000 years ago), and LGM (22,000 years ago). Bioclimatic variables were then prepared for input into Maxent 3.4.4 using QGIS 3.4 (QGIS.org, 2022) following De Alban (2022). Polygon shapefiles were first created for each species, which included the present-day breeding distribution as well as areas south of that to accommodate for potential shifts in distribution around the LGM. These polygons were then used to crop the bioclimatic variable raster for each respective species (Conrad et al., 2015).

We applied Maxent 3.4.4, which makes use of presence-only data and environmental data to model species’ geographical distributions (Phillips et al., 2006). Species-specific Maxent analyses were performed using the respective breeding occurrence records, background points and present-day bioclimatic variables of each species. To reduce collinearity among predictors, we removed predictors with a high variance inflation factor (>3) for each species. To facilitate parameter tuning, 20 candidate models were built for each species and evaluated using the R package ENMeval 2.0.3, testing combinations of feature classes (L, LQ, LQH, LQPH) and regularisation multipliers (0.5, 1, 2, 3, 4) (Kass et al., 2021; Merow, Smith, & Silander, 2013). To test for model overfitting and transferability, candidate models cross-validated using the ‘block’ partitioning technique (i.e., occurrences and background points were partitioned into four spatial blocks, where occurrence numbers among partitions are equal) (Fourcade, Besnard, & Secondi, 2018; Muscarella et al., 2014). Candidate models with omission rates (minimum training presence threshold) exceeding 0.2 were rejected. The candidate model with the highest area under the receiver-operator curve (AUC) was selected as the final model (Table S4) and used to predict suitable breeding area under present-day, mid-Holocene, and LGM climate conditions (Table S5).

Predicted species distributions were visualised in R (R Core Team, 2022). We performed a binary classification of predicted occurrence probability using the maximum sum of sensitivity plus specificity threshold (C. Liu, White, & Newell, 2013) and calculated suitable breeding area using the R package raster 3.5-15.

## Data availability

DNA reads generated in this study are available on Sequence Read Archive under BioProject PRJNA742889. Reference genome generated in this study is available on GenBank under [accession number] (in submission). Pipelines and analysis codes are available upon request.

## Acknowledgements

We thank the following personnel and institutions for their generous contribution of samples (Table S1): Paul Sweet and Thomas Trombone at the American Museum of Natural History (AMNH, New York); Robert Palmer and Leo Joseph at the Australian National Wildlife Collection (ANWC, Canberra); Molly Hagemann at the Bernice Pauahi Bishop Museum (BPBM, Hawaii); David Allan at the Durban Natural Science Museum (DNSM, Durban) and Celine Santillan who assisted in sample transport; Ben Marks at the Field Museum of Natural History (FMNH, Chicago); Foo Maosheng at the Lee Kong Chian Natural History Museum (LKCNHM, Singapore); Carla Marangoni and Gloria Svampa at the Museo Civico di Zoologia (MCZR, Rome); Henry McGhie at the University of Manchester, Manchester Museum (MMUM, Manchester); Robert Prŷs-Jones, Mark Adams, Alex Bond, Ari Benucci and Douglas Russell at the Natural History Museum, London (NHMUK, Tring); Manuel Schweizer at the Naturhistorisches Museum der Bürgergemeinde Bern (NMBE, Bern); Bob McGowan at the Natural Museum of Scotland (NMS, Edinburgh); Joanna Sumner at Museums Victoria (NMV, Melbourne); Angela Ross at the National Museums NI (NMNI, Northern Ireland) and David Allen and Graeme Buchanan who assisted in sample transport; Jan Bolding Kristensen at the Natural History Museum of Denmark (SNM, Copenhagen); José Alves and Camilo Carneiro at the University of Iceland (UOI, Reykjavik); Sharon Birks at the Burke Museum, University of Washington (UWBM, Seattle); Pavel S. Tomkovich, Dmitry Shitikov and Vladimir Sotnikov at the Zoological Museum of Moscow State University (ZMMU, Moscow); and Fyodor Kondrashov and Lisa Chilton who assisted in sample transport. Fletcher Smith assisted in Mozambique (with permission from Lucilia Chuquela, Museu de História Natural and Universidade Eduardo Mondlane, Maputo) with further assistance from Rebecca and Cyril Kormos, Patricia Zurita and Vinayagan Dhamarajah. HZT acknowledges Elize Ying Xin Ng, Pratibha Baveja, Yong Chee Keita Sin, Shivaram Rasu, Dominic Yong Jie Ng, Meng Yue Wu, Liu Xiaoming, and Jose Don De Alban for assistance with laboratory procedures and analyses. The authors acknowledge support from the National Genomics Infrastructure in Stockholm funded by Science for Life Laboratory, the Knut and Alice Wallenberg Foundation and the Swedish Research Council, and SNIC/Uppsala Multidisciplinary Center for Advanced Computational Science for assistance with massively parallel sequencing and access to the UPPMAX computational infrastructure. MI acknowledges support from the Swedish Research Council (2019-03900). KMG acknowledges support from the DBT-Ramalingaswami Fellowship (BT/HRD/35/02/2006). BC acknowledges funding from the South East Asian Biodiversity Genomics (SEABIG) Grant (WBS R-154-000-648-646 and WBS R-154-000-648-733) and startup funding from Trivedi School of Biosciences, Ashoka University. Finally, FER acknowledges a Singapore Ministry of Education Tier 2 grant (R-154-000-C41-112), which funded most labwork.

## Author Contributions

FER, JJFJJ, GAA, HZT conceptualised the research aims. JJFJJ, GAA, GC, and HZT collected samples, KMG and BC designed the probe set and MI constructed the reference genome. HZT performed all laboratory procedures with guidance from CYG, KMG, and BC. HZT performed all bioinformatic analyses with guidance from CYG, SEHP, and FER. HZT and FER produced the initial draft of the manuscript which was reviewed by all co-authors.

