## Supplementary information for "Megafaunal extinctions—not climate change—seem to explain Holocene genetic diversity declines in *Numenius* shorebirds"

### Supplemental Information

Table S1. Details of samples collected for this study. Species and subspecies identities are given, if available, as identified by the label information provided by the source museum or institution except American samples of *Numenius phaeopus*, which is now widely separated as *N. hudsonicus* (Gill, Donsker, & Rasmussen, 2021; Tan et al., 2019). Samples denoted with an asterisk (*) were not included in the final SNP or sequence dataset due to high missing data. Sample NBME 1039630, denoted with a caret (^), was identified as *N. tenuirostris* on its museum label but should be reassigned to *N. arquata* based on our analyses. Sequence reads are available in the Sequence Read Archive repository (BioProject number: PRJNA742889). Abbreviations: AMNH – American Museum of Natural History, New York; ANWC – Australian National Wildlife Collection, Canberra; BPBM – Bernice Pauahi Bishop Museum, Hawaii; DNSM – Durban Natural Science Museum, Durban; FMNH – Field Museum of Natural History, Chicago; GA – Gary Allport, fieldwork; LKCNHM – Lee Kong Chian Natural History Museum, Singapore; MCZR – Museo Civico di Zoologia, Rome; NHMUK – Natural History Museum, London (Tring); NMBE – Naturhistorisches Museum der Bürgergemeinde Bern, Bern; NMNI – National Museums NI, Northern Ireland; NMS – Natural Museum of Scotland, Edinburgh; NMV – Museums Victoria, Melbourne; UOI – University of Iceland, Reykjavik; UWBM – Burke Museum, University of Washington, Seattle; ZMMU – Zoological Museum of Moscow State University, Moscow.

| **Museum/ catalogue number** | **Scientific name** | **Country of collection** | **Collection locality** |
|---|---|---|---|
| NHMUK 1965.M.3011 | *Numenius americanus* | USA | Texas, Mueces Co. |
| NHMUK 1965.M.3012 | *Numenius americanus* | USA | Texas, Mueces Co. |
| UWBM 56311 | *Numenius americanus* | USA | Montana, Fort Benton |
| UWBM 89517 | *Numenius americanus* | USA | Washington, Yakima |
| AMNH DOT 20952 | *Numenius arquata arquata* | Sweden | Norrbotten, Ängesbyn |
| ZMMU 084 | *Numenius arquata suschkini* | Russia | Orenburg, Akbulak, Ilek River |
| ZMMU RYA2439 | *Numenius arquata orientalis* | Russia | Altai Krai, Uglovsky, Lyapunikha Lake |
| NHMUK 1960.12.67 | *Numenius arquata orientalis* | Maldives | Maldive Islands, Gan |
| ZMMU RYA2355 | *Numenius arquata suschkini* | Russia | Altai Krai, Blagoveshenka, Mikhailovka |
| FMNH 407302 | *Numenius arquata suschkini* (type) | Senegal | Degama |
| NHMUK 1902.8.14.1 | *Numenius arquata* | Azerbaijan | Baku, Kuba District, Lake Akzi-Birr |
| AMNH DOT 10893 | *Numenius arquata arquata* | UK | England, Lancashire, Barrow in Furness |
| ZMMU KEV013 | *Numenius arquata suschkini* | Russia | Ryazan', Korablinsky, Semion |
| MCZR 15733 | *Numenius arquata* | Italy | Veneto, Northeast Italy |
| UWBM 74389 | *Numenius arquata suschkini* | Russia | Kirovskaya Oblast, Zuyevskiy Rayon, Beregovoy Station, Tcheptsa river |
| UWBM 73571 | *Numenius arquata orientalis* | Russia | Avtonomnaya Respublika Buryatiya, Kabanskiy Rayon, Ulan-Ude, Mursino |
| ZMMU 807 | *Numenius arquata suschkini* | Russia | Orenburg |
| NHMUK 1896.7.1.780 | *Numenius borealis* | USA | Massachusetts, Ipswich |
| NHMUK 1922.3.5.558 | *Numenius borealis* | Guyana | Potaro-Siparuni |
| NHMUK 1965.M.2961 | *Numenius borealis* | USA | Texas, Brownsville |
| NMS.Z 1870.28.78 c/1 | *Numenius borealis* | Canada | Northwest Territories, Anderson River |
| *NHMUK E/1902.3.10.54 | *Numenius borealis* | Canada | Northwest Territories, Anderson River |

| | | | |
|---|---|---|---|
| ZMMU 108581 | *Numenius hudsonicus* | Canada | Manitoba, Churchill |
| ZMMU 108582 | *Numenius hudsonicus* | Canada | Quebec, Lurd-de-Blanc-Sablor |
| UWBM 53963 | *Numenius hudsonicus* | USA | Alaska, Paxson, along Denali Highway |
| UWBM 43588 | *Numenius hudsonicus* | USA | Washington, Grays Harbor, Westport, mud flats at north edge of South Bay |
| UWBM 69057 | *Numenius hudsonicus* | Nicaragua | Departamento de Rivas, La Flor |
| ANWC B34783 | *Numenius madagascariensis* | Australia | Queensland, Cairns International Airport |
| ANWC B50496 | *Numenius madagascariensis* | Australia | Mary Island North, King Sound, Northwest of Derby |
| ANWC B51518 | *Numenius madagascariensis* | Australia | Queensland, Aurukun Region, South of Weipa, Cape York Peninsula |
| ANWC B33574 | *Numenius minutus* | Australia | Northern Territory, Koolpinyah Station, East of Darwin |
| NMV Z5885 | *Numenius minutus* | Australia | Northern Territory, Cox Peninsula Rd, Berry Springs |
| NHMUK 1931.8.18.906 | *Numenius phaeopus* | Madagascar | North Madagascar, Anomontsangana |
| DNSM 1043 | *Numenius phaeopus alboaxillaris* | South Africa | KwaZulu-Natal, Durban, Durban Bay |
| NHMUK 1882.12.3.2 | *Numenius phaeopus alboaxillaris* | Kenya | Mombasa |
| ZMMU 43101 | *Numenius phaeopus alboaxillaris* | Kazakhstan | Kostanaj, Toktas Lake |
| GA | *Numenius phaeopus alboaxillaris* | Mozambique | Maputo Bay |
| NHMUK 1903.10.14.292 | *Numenius phaeopus alboaxillaris* (type) | Mozambique | Inhambane |
| ZMMU 111271 | *Numenius phaeopus alboaxillaris* | Russia | Bashkortostan, Dertyuli, Russkij Angasyak |
| ZMMU 261 | *Numenius phaeopus alboaxillaris* | Russia | Bashkortostan, Abzelilovo, Maly Kizel Lake |
| DNSM 1044 | *Numenius phaeopus alboaxillaris* | South Africa | KwaZulu-Natal, Durban, Durban Bay |
| ZMMU 73801 | *Numenius phaeopus alboaxillaris* | Turkmenistan | Balkansky, Chuikishlyar |
| NHMUK 1894.2.19.101 | *Numenius phaeopus* | - | East Africa, Fantee, Naqua River |
| UOI 581748 | *Numenius phaeopus islandicus* | Iceland | Southern Region |
| UOI 589357 | *Numenius phaeopus islandicus* | Iceland | Southern Region |
| ZMMU 953 | *Numenius phaeopus phaeopus* | Russia | Krasnoyarsk, Evenkia, Severnoe Lake |
| ZMMU 38305 | *Numenius phaeopus phaeopus* | Russia | Yamalo-Nenets, Salekhard |
| ZMMU SAP 8327 (302) | *Numenius phaeopus phaeopus* | Russia | Kaliningrad, Curonian Spit |
| ZMMU 950 | *Numenius phaeopus phaeopus* | Russia | Kirov, Podosinovets, Ul'skoe Mire |
| UWBM 49676 | *Numenius phaeopus phaeopus* | Russia | Murmanskaya Oblast, Teriberka |
| *UWBM 59485 | *Numenius phaeopus* | Russia | Yamalo-Nenetskiy Avtonomnyy Okrug, Labytnangi |
| ZMMU 94691 | *Numenius phaeopus rogachevae* | Russia | Krasnoyarsk, Evenkia, Ukikit Lake |
| ZMMU 353 | *Numenius phaeopus variegatus* | Russia | Chukotsky, Anadyr, Meinypil'gyno |
| ZMMU 952 | *Numenius phaeopus variegatus* | Russia | Chukotsky, Anadyr, Kanchalan River, 50 km N of Kanchalan Village |

| | | | |
|---|---|---|---|
| *ANWC B33977 | *Numenius phaeopus variegatus* | Australia | Northern Territory, Victoria River Mouth, North of Bullo River Homestead |
| *ANWC B51458 | *Numenius phaeopus variegatus* | Australia | Queensland, Kalpowar Station, Princess Charlotte Bay, Cape York Pen |
| BPBM 185971 | *Numenius tahitiensis* | USA | Hawaii, Oahu, James Campbell NWR, Ki'I kiosk |
| BPBM 186246 | *Numenius tahitiensis* | USA | Hawaii, Kauai |
| *AMNH DOT 10919 | *Numenius tahitiensis* | USA | Hawaii, Honolulu, French Frigate Shoals, Tern Island, Hawaiian Islands NWR |
| NHMUK 1918.1.10.1 | *Numenius tenuirostris* | Palestine | South Palestine |
| NHMUK 1933.2.16.151 | *Numenius tenuirostris* | Iraq | Tigris, Amara river |
| MCZR 15761 | *Numenius tenuirostris* | Italy | Lucca (Tuscany) |
| NMNI | *Numenius tenuirostris* | Italy | - |
| NMNI | *Numenius tenuirostris* | - | - |
| ^NBME 1039630 | *Numenius tenuirostris* | Russia | Siberia |
| LKCNHM 103168 | *Tringa totanus* | Singapore | Sungei Buloh Wetlands Reserve |

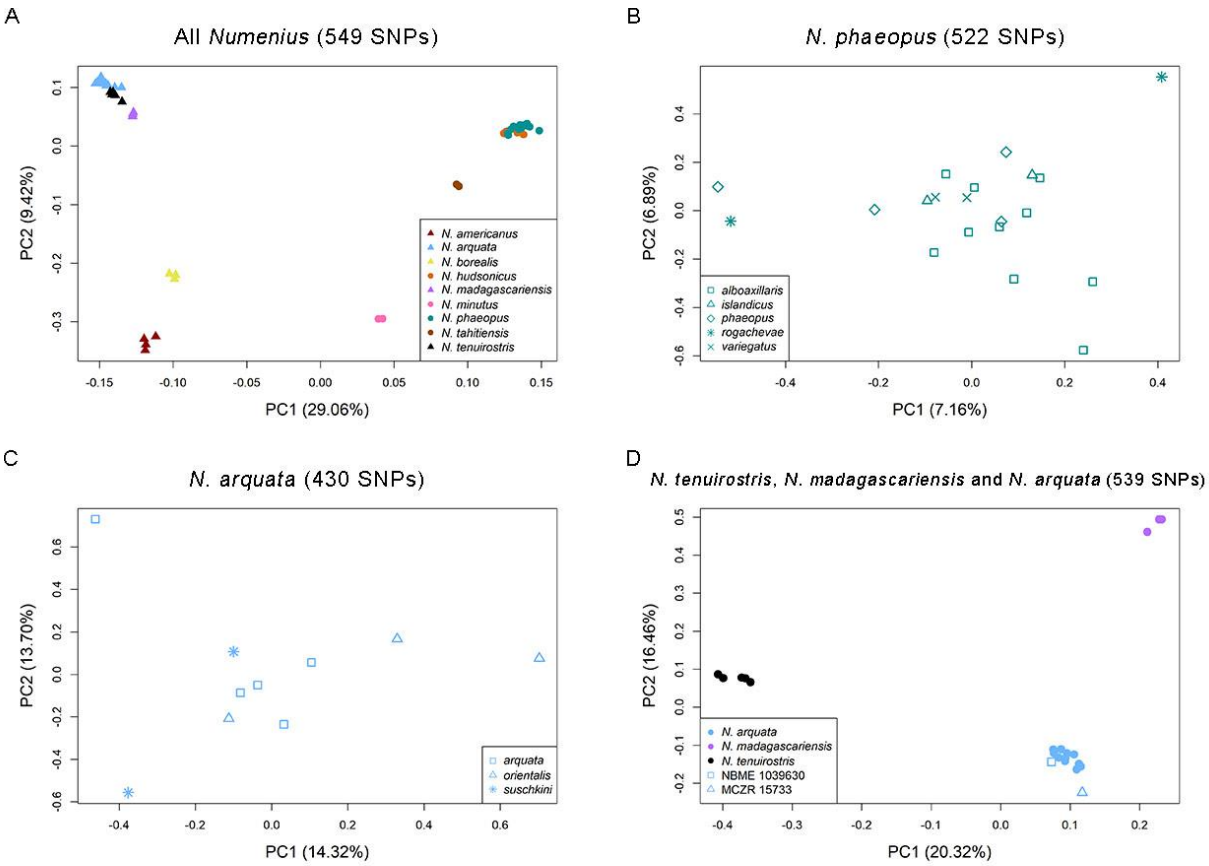


Figure S1. Principal component (PC) analysis of *Numenius* samples, with percentage of variation of the two most important PCs displayed. Colours correspond to species identities in figure 1. The four plots display PC analyses of (A) all *Numenius* species (B) Eurasian whimbrel *N. phaeopus* (C) Eurasian curlew *N. arquata* and (D) Palaearctic curlews *N. tenuirostris*, *N. madagascariensis* and *N. arquata*.

Table S2. Evolutionary distinctness, phylogenetic diversity and evolutionarily distinct and globally endangered (EDGE) scores of *Numenius* species, as calculated from MCMCTree branch lengths and IUCN status (Jetz et al., 2014). IUCN (2020) status abbreviations: CR – Critically Endangered; EN – Endangered; NT – Near Threatened; LC – Least Concern. Asterisks (*) given for putatively extinct species.

<table>
<tr><th>Clade</th><th>Species</th><th>IUCN Status</th><th>Evolutionary distinctness in million years (MY)</th><th colspan="2">Phylogenetic diversity (my)</th><th>EDGE score</th></tr>
<tr><td rowspan="4">whimbrels</td><td>N. tahitiensis</td><td>NT</td><td>3.18</td><td rowspan="4">12.23</td><td rowspan="9">25.10</td><td>2.12</td></tr>
<tr><td>N. phaeopus</td><td>LC</td><td>2.59</td><td>1.28</td></tr>
<tr><td>N. hudsonicus</td><td>LC</td><td>2.59</td><td>1.28</td></tr>
<tr><td>N. minutus</td><td>LC</td><td>3.87</td><td>1.58</td></tr>
<tr><td rowspan="5">curlews</td><td>N. tenuirostris</td><td>CR*</td><td>2.31</td><td rowspan="5">12.87</td><td>3.97</td></tr>
<tr><td>N. arquata</td><td>NT</td><td>2.31</td><td>1.89</td></tr>
<tr><td>N. madagascariensis</td><td>EN</td><td>2.43</td><td>3.31</td></tr>
<tr><td>N. americanus</td><td>LC</td><td>2.91</td><td>1.36</td></tr>
<tr><td>N. borealis</td><td>CR*</td><td>2.91</td><td>4.14</td></tr>
</table>

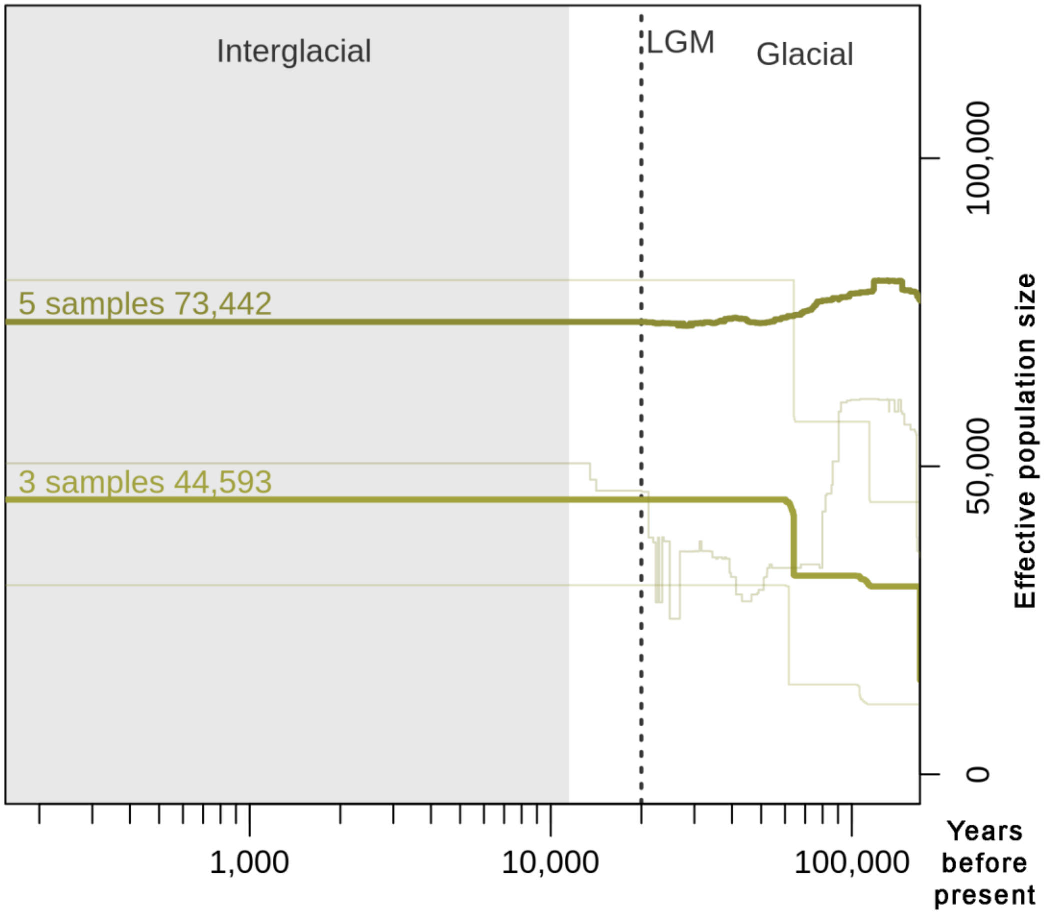


Figure S2. Demographic history reconstruction using stairway plot for *N. borealis*, showing results for two datasets, one containing all five samples and the other being a subset of three samples with low missingness. Present-day effective population size of each dataset is indicated above the lines.

Table S3. Details of occurrence points in breeding areas used for input into Maxent for four *Numenius* species, including sample size, month (x denotes the month for which records are included), year and sources of records.

| **Species** | **Number of samples** | **Month** | | | | **Year** | **Source** |
|---|---|---|---|---|---|---|---|
| | | **Apr** | **May** | **Jun** | **Jul** | | |
| *N. phaeopus* | 158 | | x | x | x | All years | eBird, 2021; GBIF.org, 2022a; Lappo, Tomkovich, & Syroeckovskiy, 2012 |
| *N. hudsonicus* | 1713 | | x | x | x | All years | eBird, 2021; GBIF.org, 2022b, 2022c, 2022d |
| *N. americanus* | 644 | x | x | x | | 1960 – 1990 | eBird, 2021; GBIF.org, 2022e |
| *N. arquata* | 997 | x | x | x | x | All years | eBird, 2021; GBIF.org, 2022f |

Table S4. Summary of the parameters and results of the best ecological niche model identified for each of the four *Numenius* target species.

| **Species** | **Feature class(es)** | **Regularisation multiplier** | **Mean validation area under the receiver operating curve (auc.val.avg)** | **Mean validation continuous Boyce index (cbi.val.avg)** | **Mean minimum training presence omission rate (or.mtp.avg)** | **Maximum test sensitivity plus specificity Cloglog threshold** |
|---|---|---|---|---|---|---|
| *N. phaeopus* | LQHP | 4 | 0.831 | 0.836 | 0.006 | 0.570 |
| *N. hudsonicus* | LQH | 2 | 0.829 | 0.691 | 0.044 | 0.408 |
| *N. americanus* | LQ | 4 | 0.881 | 0.733 | 0.002 | 0.459 |
| *N. arquata* | L | 0.5 | 0.827 | 0.757 | 0.002 | 0.477 |

Table S5.Visualisation of ecological niche model results with green corresponding to a higher probability of presence and brown corresponding to a lower probability. Numbers below each plot represents the total suitable breeding area for each study species and the respective times.

| **Species** | **1960-1990** | **mid-Holocene (6,000 years ago)** | **Last Glacial Maximum (22,000 years ago)** |
|---|---|---|---|
| *N. phaeopus* | 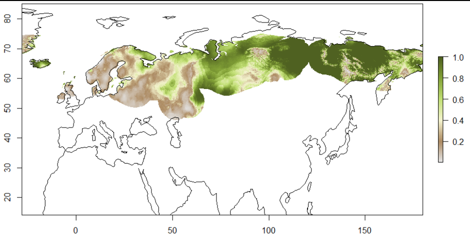 7,419,464.0 km$^2$ | 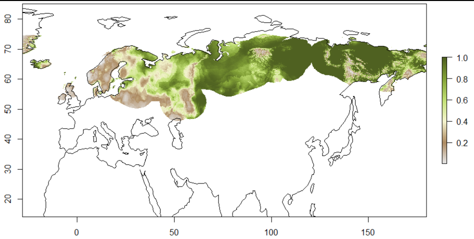 9,203,730.0 km$^2$ | 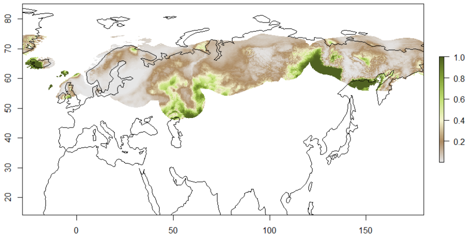 1,794,131.0 km$^2$ |
| *N. hudsonicus* | 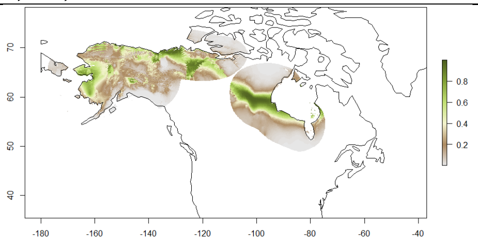 1,187,597.0 km$^2$ | 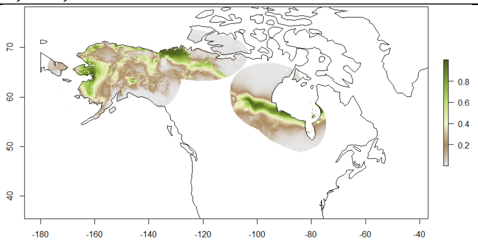 981,343.5 km$^2$ | 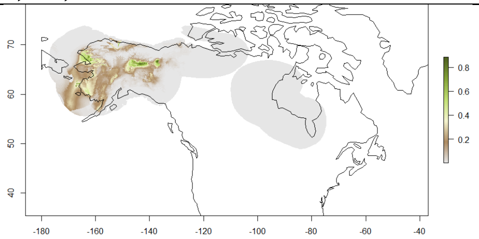 116,941.3 km$^2$ |
| *N. americanus* | 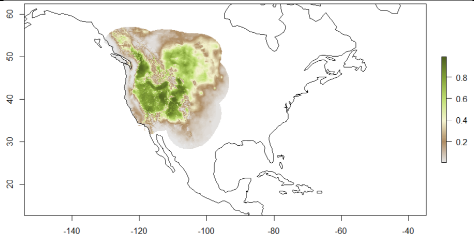 1,935,383.0 km$^2$ | 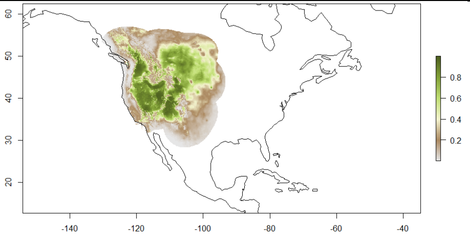 2,399,962.0 km$^2$ | 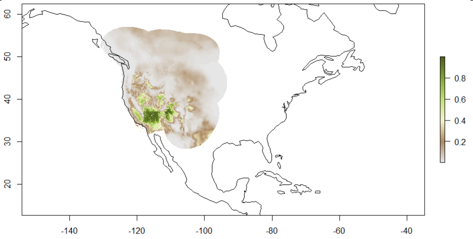 352,982.3 km$^2$ |
| *N. arquata* | 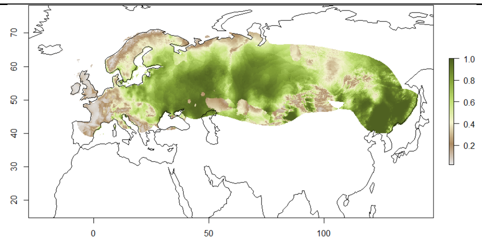 14,466,715.0 km$^2$ | 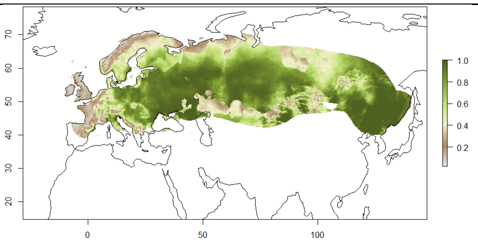 16,143,041.0 km$^2$ | 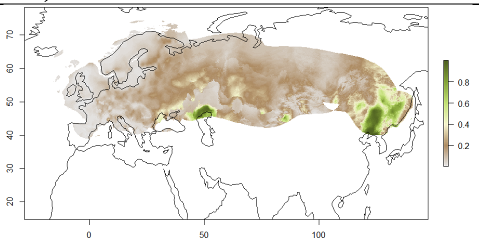 1,295,521.0 km$^2$ |
